# Connectomic traces of Hebbian plasticity in the entorhinal-hippocampal system

**DOI:** 10.1101/2025.04.07.647631

**Authors:** Eleonora Grasso, Maria Corteze, Helene Schmidt

## Abstract

The key model of how we learn and memorize is Hebbian learning in the hippocampus, via long-term potentiation of synapses, allowing the storage of associations, linkage to places, and their consolidation into imprinted episodes. Learning is therefore expected to change synaptic weights. With the notion of hippocampal circuits being the primary site of learning in the mammalian brain, it has been assumed that all synapses in these circuits are constantly exposed to synaptic plasticity, and possibly in a learned state. However, a testing of these hypotheses is so far missing. In particular, the systematic mapping of synaptic weight distributions, and their relation to Hebbian preconditions has not been achieved yet in the hippocampal-entorhinal system. Here, we report such a systematic connectomic mapping of synaptic weight distributions and their relation to same-axon same-dendrite paired synaptic configurations across the hippocampal-entorhinal system. By analyzing millions of synapses and tens of thousands of paired synaptic configurations from 3D EM-based automated circuit reconstructions in hippocampal areas CA3, CA1, and layers 2 and 3 of the medial entorhinal cortex (MEC), we found systematic and unique synaptic weight distributions, with almost 50% (but not 100%) of synaptic weights in CA1 being in a Hebbian-consistent state, CA3 uniquely exhibiting small synaptic weights with indications of learned states, and MEC resembling previous data from other isocortices, with only up to 20% learned synaptic configurations. We further analyzed the sublayer-specificity of these weight distributions, finding the molecular layer of CA1 and the lower layers of CA3 being the most unique site of potential learning. Together, this data provides a first systematic synaptic weight analysis of the key neuronal system involved in memory formation in mammalian brains.

## INTRODUCTION

In today’s artificial intelligence, storage of information in the (artificially modelled) synaptic weights of large neuronal networks is standard. This has been strongly inspired by neuroscience’s models of learning via synaptic creation, strengthening (and weakening and abolition). In particular the hippocampus has been studied extensively as a memory structure, where long-term potentiation is thought to underly the storage of memory traces. But while in artificial intelligence, analysis of synaptic weight matrices and their storage properties is directly accessible, for the mammalian memory system, similar data is not yet available.

With the possibilities of 3D EM-based connectomic mapping, however, systematic analysis of synaptic weights in these crucial circuits has become more realistic. In fact, pioneering studies (Bartol, Bromer et al. 2015, Samavat, Bartol et al. 2024) have analyzed even small volumes of EM data from the hippocampus and conducted spearheading analyses about information storage capacity, traces of LTP and effects of strong stimulation.

Yet, basic assumptions of our understanding of the hippocampal-entorhinal system remain untested. In particular, it is not known what fraction of synapses in CA1, CA3 and other areas are at any given point in time in a state resulting from learned (LTP) configurations. And is storage graded, or binary (Dorkenwald, Turner et al. 2022)? Are the same synaptic weight ranges and principles used in the various parts of the circuit? Are there specific hot-spots of synaptic imprinting (Sievers, Motta et al. 2024)? Is storage primarily in the local circuit, or does it involve long-range connections (Kitamura, Pignatelli et al. 2014, Uytiepo, Zhu et al. 2025)? And how is the MEC, the key partner structure, involved?

All of these questions could be addressed from a systematic connectomic mapping of the relevant structures using 3D EM based analysis. Here, we provide such data. For our analyses, we specifically avoided any “stimulation” or “learning” paradigm - we wanted to analyze the system in a realistic (if laboratory-environment based) setting. For this, brains of 4-week old mice were analyzed using state-of-the art connectomic methods, providing glimpses at the synaptic network thought to underly mammalian memory formation.

## RESULTS

We used our previously published large-scale 3D EM data from mouse CA3 (Sammons, Vezir et al. 2024), CA1 (Corteze, Grasso et al. 2025) and two cubes of 3D EM data from mouse MEC for automated segmentation and synapse detection, yielding a total of 5.85 million synapses for connectomic analysis. Using previously established methodology relating synaptic size (axon-spine interface size, (Cheetham, Barnes et al. 2014, de Vivo, Bellesi et al. 2017, Motta, Berning et al. 2019)) to synaptic strength, and studying same axon-same dendrite paired connections in comparison to random pairs of synapses (Bartol, Bromer et al. 2015), we analyzed the distributions of synaptic weights and their relation to a possible history of Hebbian plasticity.

### Overall synaptic size distributions

First, we mapped the synaptic size distributions using all spine synapses automatically detected in cubes of 3D EM data (Fig. 1) from CA1 (n=1,666,113 synapses), CA3 (n= 3,448,083 synapses), and MEC layer 2 (n=275,908 synapses) and layer 3 (n=464,091 synapses). The overall synaptic size distributions showed already starkly different profiles (Fig. 1B-E): while the data from MEC (Fig. 1D,E) was very similar to other data from neocortex (Motta, Berning et al. 2019) with an almost log-normal shape (with a slant towards larger connections), CA1 (Fig. 1B) and CA3 (Fig. 1C) showed highly structured, non-log normal distributions: in CA1, a clear double peak, possibly from at least two underlying size distributions, was visible, with smaller and larger synapses at comparable prevalence (Fig. 1B). In CA3, however, small synapses far outnumbered the larger synaptic distributions (by almost 9-to-1), but a “shoulder” of larger synapses was clearly detectable (Fig. 1C). Notably, the synaptic size peaks between CA1 and CA3 were clearly matched (Fig. 1F), and differed from the main synaptic sizes found in MEC. These findings triggered important questions: were the over-abundant small synapses in CA3 just randomly established, or could systematic patterns of innervation be found? Could we detect an overabundance of paired synaptic configurations consistent with a history of Hebbian plasticity? What fraction of the synapses in CA1 (and CA3) would be consistent with a Hebbian state? And was any of this dependent on the layer location within these brain areas?

**Figure 1.**
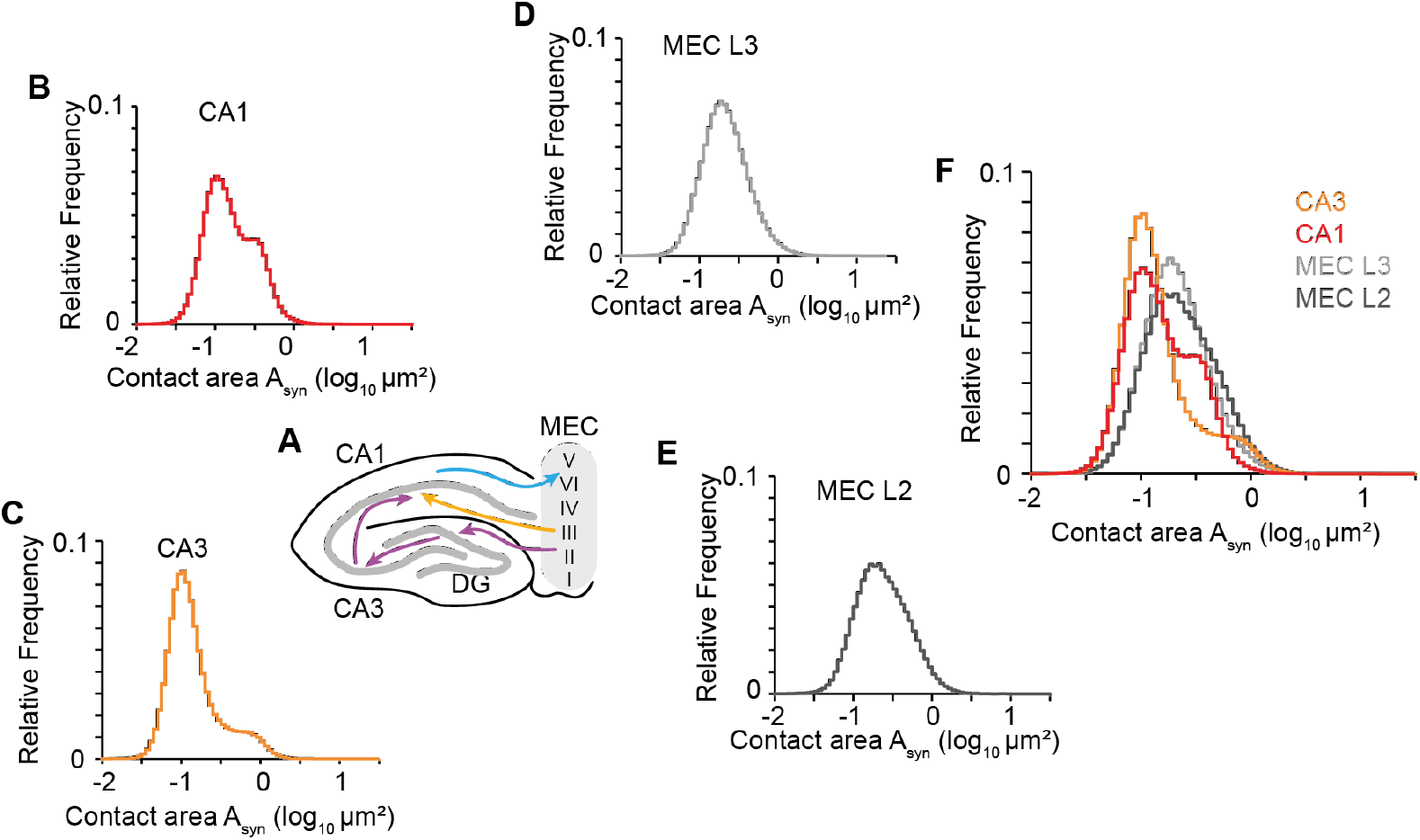
Analysis of synaptic size distributions in the hippocampal-entorhinal system. **(A)** Sketch of most of the so-far known circuits between medial entorhinal cortex (MEC) with its distinct layers, and the hippocampus with areas cornu ammonis 1 (CA1) and CA3. From MEC layers 2 and 3, and CA1 and CA3, 3D EM datasets were analyzed to obtain a systematic mapping of the synaptic size distributions. **(B-D)** Synaptic size distributions from all spine synapses in subvolumes of CA1, CA3, MEC L2 and L3, respectively. Note succinct shapes of these distributions. **(E)** Overlay of synaptic size distributions shows unique populations of small and large synapses in hippocampus, but not MEC, that are precisely size matched between CA3 and CA1.

### Analysis of same-axon same-dendrite synaptic pairs

We detected pairs of synapses made by the same axon with the same dendrite (SASD) and compared the synaptic sizes in these pairs to those of synapse pairs randomly drawn from the dataset (Fig. 2, (Sorra and Harris 1993, Bartol, Bromer et al. 2015, Bloss, Cembrowski et al. 2018, Motta, Berning et al. 2019)). Synapses drawn from SASD pairs were slightly larger than the overall distribution, consistent with a view of synaptic pairs being formed to further strengthen synaptic connections beyond the weight of a single synapse. This effect was substantial in MEC L2 (Fig. 2B), but only modest in the other regions. Then, we compared the synaptic size difference in SASD pairs vs. randomly drawn synaptic pairs (Fig. 2C). An overabundance of low-variability synaptic pairs was clearly found in CA1 and CA3, and less so in MEC L2 and L3. Notably, even in CA1, considered to be the key site of synaptic plasticity in the mammalian brain, only about half of the synaptic configurations showed an overabundance of low-variability synaptic pairs (compare blue vs black in Fig. 2C), inconsistent with a view of CA1 as completely governed by saturated LTP, at least in the non-stimulated animal. When analyzing in more detail the dependence of synaptic size similarity for smaller vs. larger synaptic pairs (Fig. 2D), we noted that an over-abundance of similar synaptic pairs was also found for the smaller synaptic weights. In particular for CA3, where the smaller synapses were so frequent, this could imply that these are not purely randomly wired, but subject to Hebbian plasticity with a small synaptic weight target state, as well.

**Figure 2.**
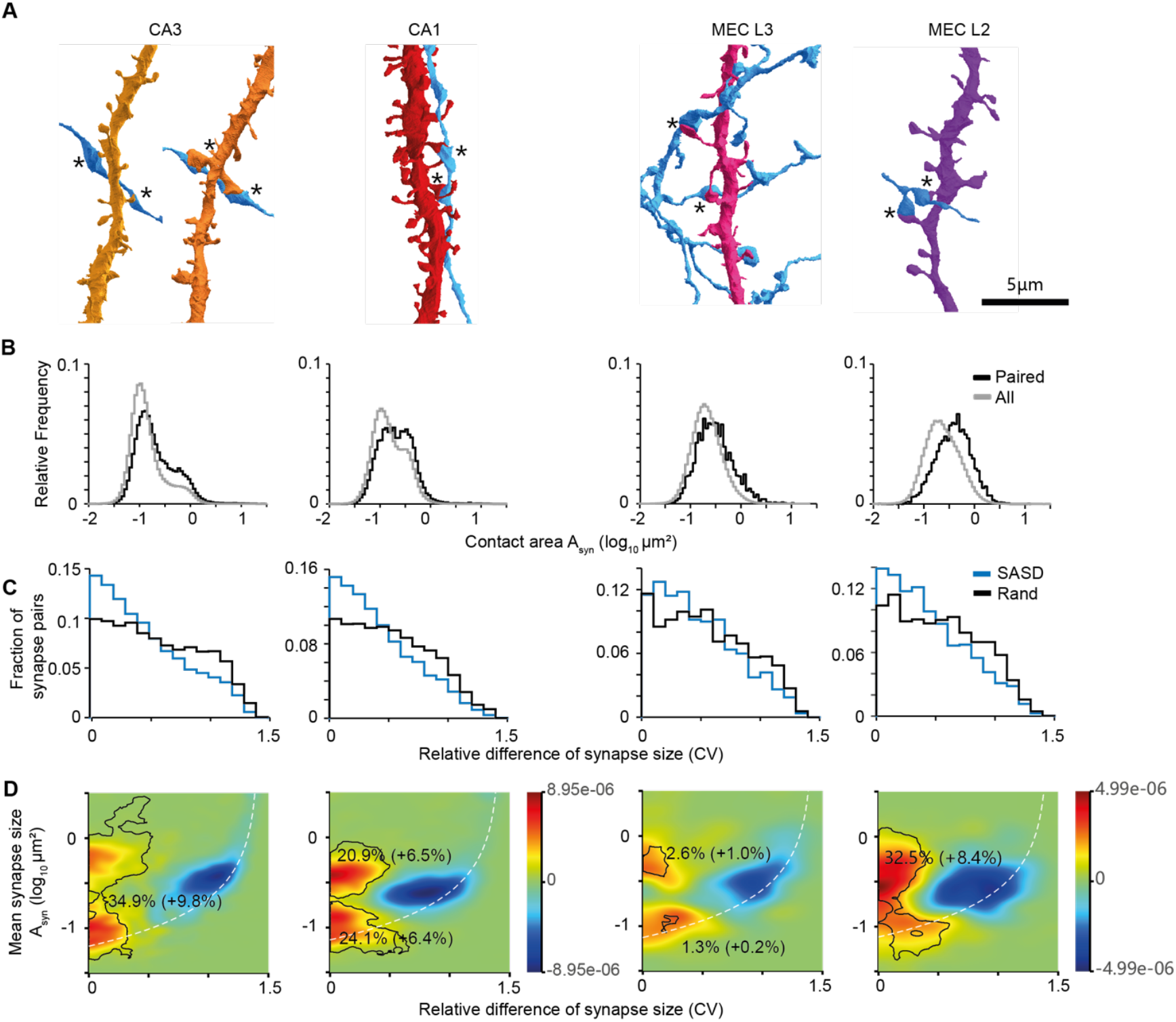
Analysis of paired synaptic configurations in same-axon same-dendrite connections for detecting possible traces of Hebbian plasticity. **(A)** Examples of dendrites and axons in CA3, CA1, MEC L2, MEC L3, respectively, that form paired synaptic innervations (same-axon same-dendrite (SASD) synapse pairs). **(B)** Synaptic sizes in SASD pairs (black), compared to the overall synaptic size distributions (grey). Note slight shift to larger synapses (most pronounced in MEC L2), and signs of bimodality in hippocampus, but not MEC. **(C)** Analysis of synaptic size variance in SASD synaptic pairs (blue) compared to randomly drawn synaptic pairs (black). Note substantial shift towards low-variance pairs in particular in the hippocampus. **(D)** Analysis of variance-size relationship indicates substantial overabundance of large and low-variance SASD pairs in CA1 and CA3, and MEC L2, but less so in MEC L3. Color scale, relative over (warm colors) or underabundance (cold colors). Isocontours indicate regions of significant overabundance of SASD vs. randomly drawn synapse pairs, numbers report fraction of synapse pairs residing within these significance regions. Analysis similar to Motta et al., 2019.

### Layer-specificity of synaptic weights in CA3

We next investigated whether synaptic weight distributions were dependent on the location across the layers of the hippocampus. In CA3 (Fig. 3), local connectivity occurs primarily via synapses located in stratum radiatum (SR) and stratum oriens (SO) (Sammons, Vezir et al. 2024), while stratum lacunosum-moleculare (SLM) is expected to comprise long-range input axons from entorhinal cortex L2. We therefore chose subvolumes from these layers in CA3 (Fig. 3A) and analyzed the layer-specific synaptic size distributions, and their relation to same-axon same-dendrite paired synapse configurations (Fig. 3B-D). All layers showed strong deviation from a log-normal synaptic size distribution (Fig. 3C). The substantial overabundance of small synapses, a unique feature of CA3 in the comparison across regions (Fig. 1), was primarily found in SR and SO, and less so in the superficial SLM (Fig. 3C). Analysis of synaptic weight variance in SASD pairs indicated an overabundance of low-variance synaptic pairs in particular for SR and SO (Fig. 3C, bottom). It showed that the population of small synapses also had significant over-similarity in SASD pairs, a possible trace of Hebbian set points for small synapses (Fig. 3D).

**Figure 3.**
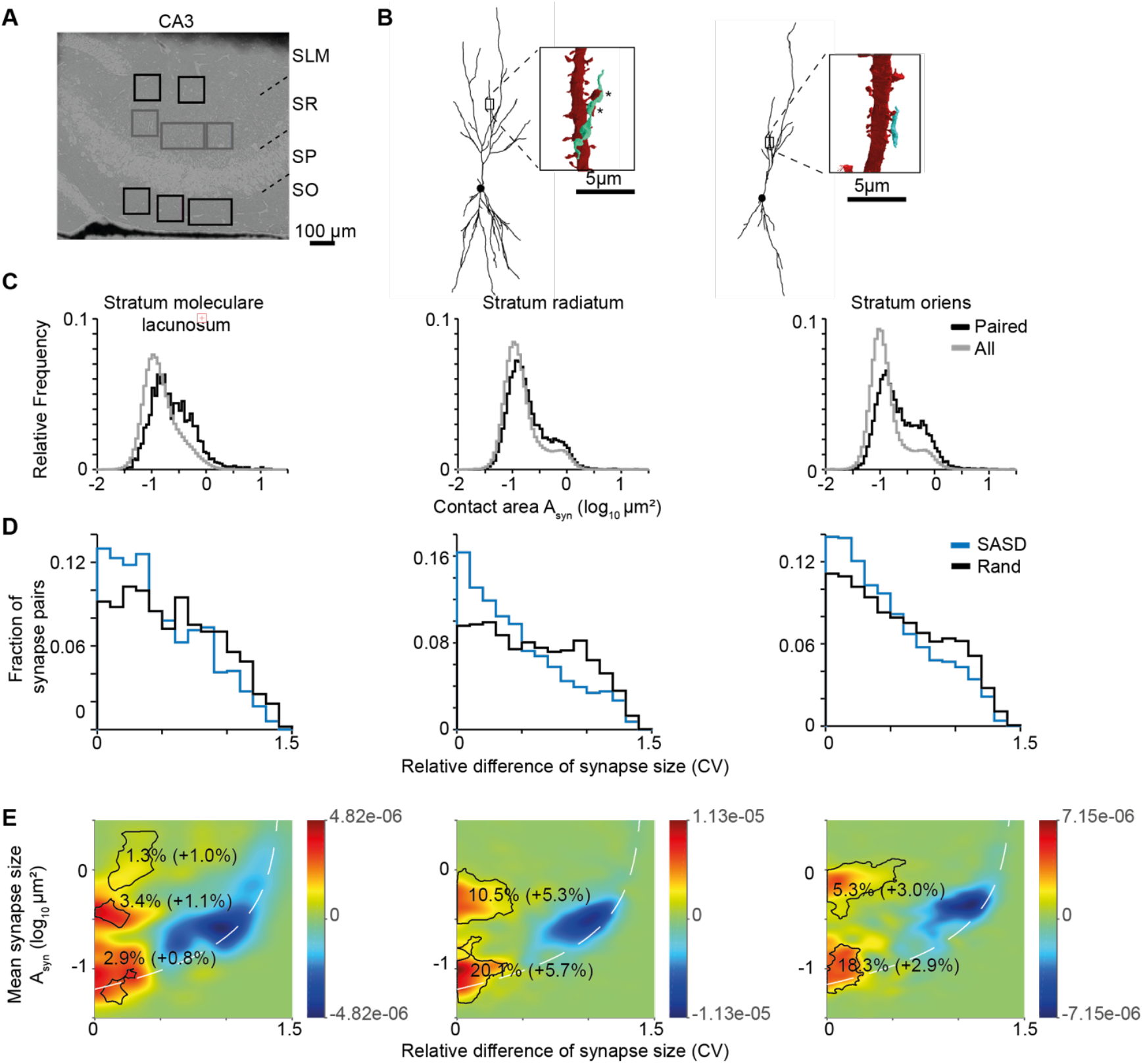
Sublayer specificity of synaptic size distributions in CA3. **(A)** Indication of bounding boxes in stratum lacunosum-moleculare (SLM), stratum radiatum (SR) and stratum oriens (SO), used for the following analyses. Note that the data in Figs. 1 and 2 was pooled from all of these bounding boxes. **(B)** Examples of same-axon same-dendrite pairs onto identified pyramidal cells in CA3 showing large and similar (left) and small and similar (right) example SASD configurations. **(C)** Over-abundance of small synapses pronounced in SR and SO, but almost abolished in SLM when considering SASD paired synaptic configurations, indicating the small-dominated circuitry to be focused in deeper CA3 layers, where most of the local (CA3-CA3) synaptic connectivity is expected. **(D**,**E)** Both 1-dimensional variance histograms and 2-d size-variance maps show pronounced traces of Hebbian-consistent low-variance SASD pairs. Note strong contribution of small-synapse but low-c.v. pairs in particular in SR and SO. E: Color scale, relative over (warm colors) or underabundance (cold colors). Isocontours indicate regions of significant overabundance of SASD vs. randomly drawn synapse pairs, numbers report fraction of synapse pairs residing within these significance regions.

### Layer-specificity of synaptic weights in CA1

We finally investigated the layer-dependency of synaptic weight distributions in CA1 (Fig. 4). Here, the overall comparison by regions had shown a substantial fraction of *large* synaptic weights with low-variance large SASD pairs (Figs. 1,2). When analyzing subvolumes in stratum lacunosum (SL), SR and SO of CA1 (Fig. 4A), we found synapse size distributions strongly deviating from log-normal shape with substantial “shoulders” of large synapses. The by far most pronounced occurrence of large synapses was in SL, with these large synapses outnumbering smaller ones in SASD pairs (Fig. 4B), a bias not found so far in any other studied cortical circuit. In all layers, a substantial fraction of low-variance SASD pairs was found (Fig. 4C) with again the most pronounced effects in SL (Figs. 4C,D).

**Figure 4.**
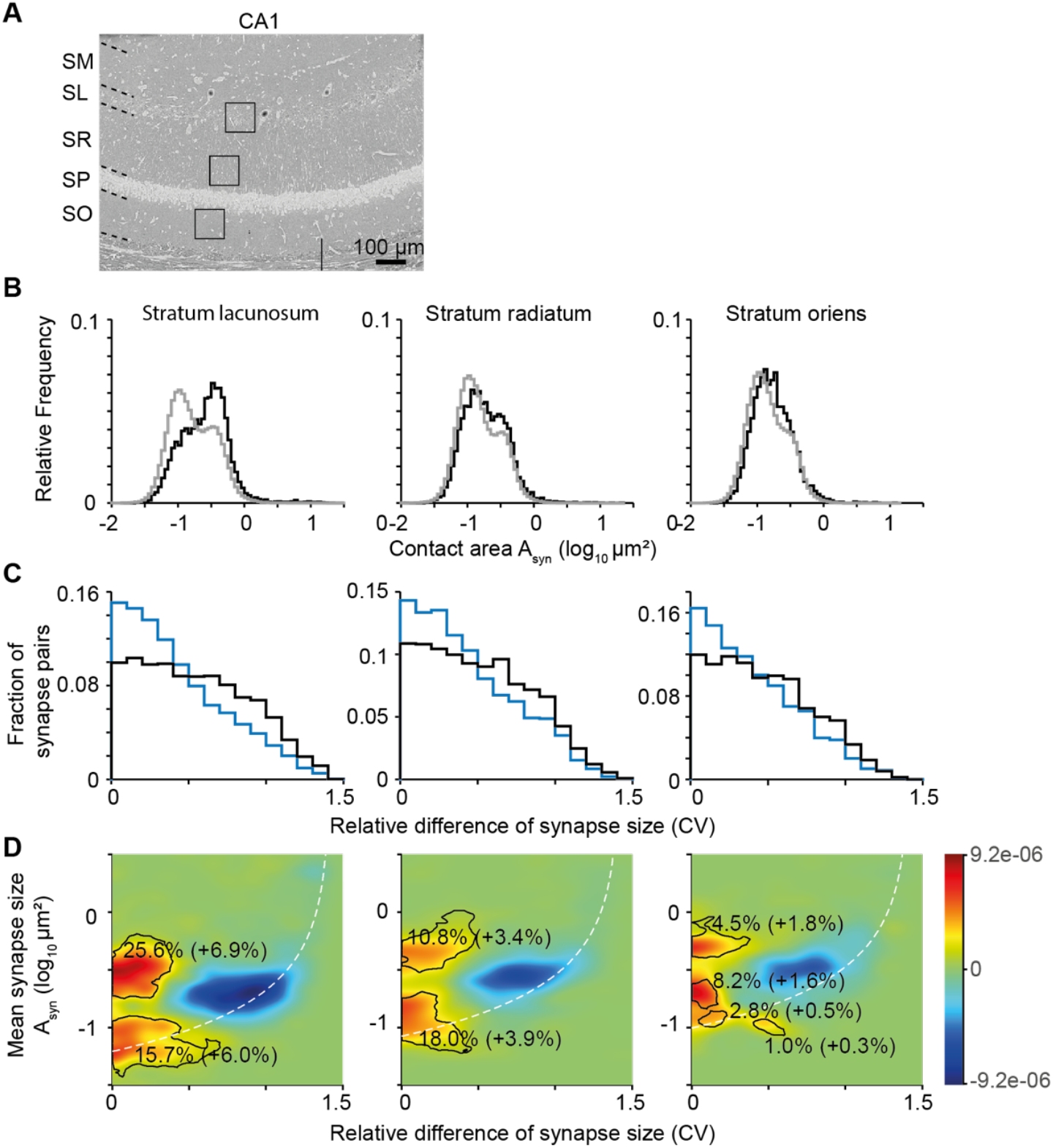
Sublayer specificity of synaptic size distributions in CA1. **(A)** Indication of bounding boxes in stratum lacunosum-moleculare (SLM), stratum radiatum (SR) and stratum oriens (SO), used for the following analyses. Note that the data in Figs. 1 and 2 was pooled from all of these bounding boxes. **(B)** Bimodal synaptic size distributions found primarily in SL of CA1, where long-range input from MEC is thought to arrive. “Shoulder” of large synapses less pronounced in SO. **(C**,**D)** Both 1-dimensional variance histograms and 2-d size-variance maps show pronounced traces of Hebbian-consistent low-variance SASD pairs, in particular for large synaptic pairs in SL. D: Color scale, relative over (warm colors) or underabundance (cold colors). Isocontours indicate regions of significant overabundance of SASD vs. randomly drawn synapse pairs, numbers report fraction of synapse pairs residing within these significance regions.

## DISCUSSION

Using 3D EM datasets from mouse hippocampus and medial entorhinal cortex, we provide a first systematic mapping of synaptic weight distributions and their possible relation to a history of Hebbian learning in the most important memory system of the mammalian brain: the hippocampal-entorhinal formation. We find unique signatures of synaptic properties: (1) The medial entorhinal cortex shows synaptic weight distributions and Hebbian-plasticity-related signatures very much in line with reports from other neocortical data; (2) CA1 and CA3 show uniquely different synaptic weight distributions: both share a population of substantially smaller synapses and a set of large synapses, which are matched in size between the hippocampal areas; (3) CA3 has an extreme over-abundance of the small synapses, outnumbering larger ones by almost 9-to-1; (4) CA1 in contrast has almost 50% of synapses in a large-weight state; (5) CA3 small synapses show signs of Hebbian low-variance configuration in same-axon-same-dendrite pairs, suggesting non-random synaptic configurations even for the small-synapse pool, possibly for information storage in small synapses; (6) even in CA1, only up to 50% of the same-axon same-dendrite synaptic pairs show significantly reduced weight variance, consistent with saturated LTP, challenging a view in which all of CA1 is involved in information storage; (7) sublayers of CA1 and CA3 have unique properties, with SR and SO being primary sites of possible Hebbian plasticity via small synapses in CA3, but SL dominating large-synapse pair consistency in CA1. Together, this is our first detailed view of the synaptic setup of the key mammalian memory system, allowing us to obtain a synapse-level understanding of mid-term information storage in the mammalian brain.

### Synaptic weight distributions

Synaptic weights in neuronal circuits have long been assumed to be log-normally distributed, and many analyses of the synaptic weights in hippocampal circuits have also used log-normal assumptions (de Vivo, Bellesi et al. 2017, Santuy, Tomas-Roca et al. 2020, Uytiepo, Zhu et al. 2025). Our data shows clearly that none of the relevant synaptic weights are log-normally distributed, especially in CA1 and CA3 (Figs. 1,3,4). Rather, unique features are discernible, with a high-prevalence small-synapse distribution in CA3 and substantial bimodal configuration in CA1. Notably, the small-synapse population in hippocampus was not found in MEC (or previous data from other cortices, (Motta, Berning et al. 2019, Dorkenwald, Turner et al. 2022)). Our data suggests that these are not merely random synapses-in-waiting, but may themselves serve information storage in consistent high-multiplicity connections.

### Analysis of possible Hebbian learning traces from paired synaptic configurations

The approach to use paired synaptic configurations for the analysis of properties of Hebbian learning in synaptic circuits has been pioneered by Seijnowski, Harris and co-workers (Bartol, Bromer et al. 2015). Originally used to study properties of Hebbian LTP from dozens of same-axon-same-dendrite configurations, in particular estimates of storage capacity (“bit depth”) per synapse (Bartol, Bromer et al. 2015), a first large-scale 3D EM analysis of the somatosensory cortex (Motta, Berning et al. 2019) on thousands of such synaptic configurations extended this approach to determine which fraction of a synaptic circuit was consistent with a history of (saturated) Hebbian LTP. In S1 cortex, only up to 20% of the synapses could be considered Hebbian (see (Sievers, Motta et al. 2024) for even lower estimates, and (Dorkenwald, Turner et al. 2022) for an alternative view of synaptic binary storage in neocortex). These data from neocortex made it plausible that in hippocampus, in particular CA1, the assumption of the entire synaptic circuit partaking in LTP-based information storage was still valid (this assumption was also the basis for the information content analysis in (Bartol, Bromer et al. 2015)). Here, we however find that even in CA1, only up to 50% of the paired synaptic configurations are consistent with a history of saturated LTP (Fig. 2,4). This raises the possibility that even in a circuit thought to be dedicated to information storage, a substantial fraction of the synapses are required for non-storage related purposes, for example regulating excitatory-inhibitory balance and maintaining stability.

### Stimulated vs. unstimulated conditions

One key difference to many previous studies of hippocampal synaptic properties and plasticity is the lack of a “stimulation” or learning paradigm in this study. Mouse brains were analyzed at 4 weeks of age, with a learning history of a typical lab mouse. On the one hand this is a disadvantage: laboratory mice lead comparatively monotonous lives, lacking many of the (existential) stimulations of their conspecifics in the wild. It is possible that with enhanced (relative to laboratory conditions) stimulation from a more enriched environment, the degree of plasticity-consistent synaptic properties could increase, and the synaptic weight distributions be even more pronounced. At the same time, the notion of studying the brain’s memory circuits in a native state, i.e. the circuit having had to store relevant information at a normal rate, without specific trained skills may be an advantage. It is certainly a baseline, and the fact that already under these conditions, clear and succinct features of synaptic weight distributions can be found, may justify this approach.

### Key sites of learning

The sublayer-specific analysis of synaptic configurations allowed us to determine possible key sites of Hebbian plasticity in the hippocampal formation: in CA3, SR and SO stood out for possible storage in small-synapse configurations (Fig. 3), and SL in CA1 for strong over-abundance of large synaptic pairs with low weight variance (Fig. 4). Such first steps may allow us to focus in on key synaptic connections involved in information storage, and thereby refine future experiments aimed at uncovering synaptic change in response to novel stimuli.

### Graded vs binary synaptic weights

One surprising result from our data is the strongly bimodal synaptic weight distribution in the hippocampus, in particular in CA1 (Figs. 1,4). While a large fraction of the literature on synaptic plasticity has implicitly or explicitly (Bartol, Bromer et al. 2015) assumed that LTP leads to synapses reaching graded synaptic weight set points, and thus synaptic storage capacity would depend on the number of such differentiable set points, recent analyses have revived the notion of binary synaptic storage, where a synapse is either in a learned or not-learned state (Dorkenwald, Turner et al. 2022). The strongly bimodal distributions in CA1 (and CA3) may suggest such a binary interpretation. However, our data suggests that also the small synaptic weights could be the result of plasticity, thus at least two learned states would be conceivable. Still, our data does not rule out a graded storage model.

### Detailed models of synaptic information storage at the network level

So far, we interpreted our data only in the context of “learned” vs. “not learned” synapses - with the notion that Hebbian LTP could lead to saturated synaptic weights that are then maintained for a certain amount of time. The synaptic configurations that are not consistent with such a saturated set point state could however also be interpreted to be in the process of forming such storage, or in the course of dissolving previously stored information. Especially in the hippocampus, which is assumed to participate only in intermediate-term information storage, a substantial turnover of synaptic weights seems likely. While the snapshot nature of 3DEM circuit data does not allow for a direct observation of plasticity events over time, more complex kinetic models of synaptic plasticity (and turnover) can however still be fitted to this dense connectomic synaptic data. With this, more detailed plasticity models can be refined and tested, and our understanding of mammalian memory formation could be substantially advanced.

## METHODS

### Animal experiments

All experimental procedures were performed in accordance with the law of animal experimentation issued by the German Federal Government under the supervision of local ethics committees, approved by the Regierungspräsidium Darmstadt, AZ: F126/1028, in compliance with the guidelines of the Max Planck Society.

### 3D EM datasets

The 3D EM data used in this study were subvolumes of larger 3D EM datasets: for CA3, the dataset published in (Sammons, Vezir et al. 2024) was used for obtaining 9 subvolumes sized 175 × 100 × 57 µm^3^ or 100 × 100 × 50 µm^3^, each. For CA1, the dataset published in (Corteze, Grasso et al. 2025) was used for obtaining 3 volume sized 100 × 100 × 50 µm^3^ each. For MEC L2 and L3, so-far unpublished data was used of 3D EM volumes centered on layer 2 and layer 3, respectively, each sized 175 × 100 × 57 µm^3^. The experimental methods were similar for all data and are described in detail in (Sammons, Vezir et al. 2024) and (Corteze, Grasso et al. 2025).

### Image analysis, synapse detection

3D EM data were aligned as described previously (Corteze, Grasso et al. 2025) and volume segmented using voxelytics (scalable minds, Potsdam). The results of the voxelytics analysis were volume objects (“agglomerates”) with type probabilities to be axon, dendrite, or other; synapse probabilities for axon-spine interfaces, and the size of these interfaces. From these data, synaptic size distributions could be directly obtained.

### Same-axon same-dendrite paired synaptic analysis

The methods for paired synapse analysis were published in (Motta, Berning et al. 2019). Briefly, synaptic pairs from same pre- and postsynaptic agglomerates were identified (same axon-same dendrite pairs, SASD), and their synaptic sizes recorded. From this, an average synaptic size of the pair as well as the “coefficient of variation”, c.v., of this synaptic pair could be computed. The latter was used as a measure of synaptic size consistency in a synaptic pair (with 0 indicating completely matched synaptic weights in a synapse pair). Random synapse pairs were drawn by shuffling the assignment within the pre- and postsynaptic population such that synaptic weight distributions were preserved, but the pre- and postsynaptic assignments were randomized.

## Author contributions

H.S. designed research; E.G., M.C. and H.S. performed research; E.G., M.C., H.S. analyzed data; H.S. wrote the paper with contributions from all authors.

The authors declare no competing interest.

## Acknowledgements

We thank the Connectomics Department at Max Planck Institute for Brain Research for support with analysis technology; in particular Sahil Loomba and Sumanth Atreya; and Daniel Werner from scalable minds for support with data segmentation.

